# Distinct Endosomal Sorting Complexes Required for Transport Components Differentially Regulate Glutamate and Gamma-Aminobutyric Acid Receptor Surface Expression

**DOI:** 10.64898/2026.06.17.732891

**Authors:** Mohamed Fouad Shalaby, Samantha Louise McLean, Sriharsha Kantamneni

## Abstract

Endosomal sorting complexes required for transport (ESCRT) regulate membrane protein trafficking through coordinated cargo selection and endosomal processing, yet their contribution to neurotransmitter receptor sorting remains to be defined. Here, we examined how modulation of distinct complex components influences the surface expression of excitatory and inhibitory neurotransmitter receptors. Using surface biotinylation and imaging approaches in heterologous cells and primary neurons, we altered tumour susceptibility gene 101 (TSG101), a core complex I component, and vacuolar protein sorting-associated protein 4A (VPS4a), an ATPase required for complex III disassembly. Reduction of tumour susceptibility gene 101 increased receptor association with early endosomes and enhanced receptor surface localisation, whereas disruption of VPS4A promoted receptor accumulation within late endosomal compartments and impaired degradative progression. Inhibitory receptor subtypes displayed variable sensitivity. Together, these findings demonstrate that endosomal sorting complex components regulate receptor surface expression through stage-specific trafficking mechanisms associated with altered receptor recycling and degradative processing.

**Graphical abstract:** Distinct ESCRT components regulate neurotransmitter receptor trafficking through stage-specific control of the endosomal pathway. ESCRT-I disruption promotes early endosomal retention and recycling, whereas ESCRT-III impairment causes late endosomal accumulation and reduced degradation, together increasing receptor surface expression (created using Biorender).

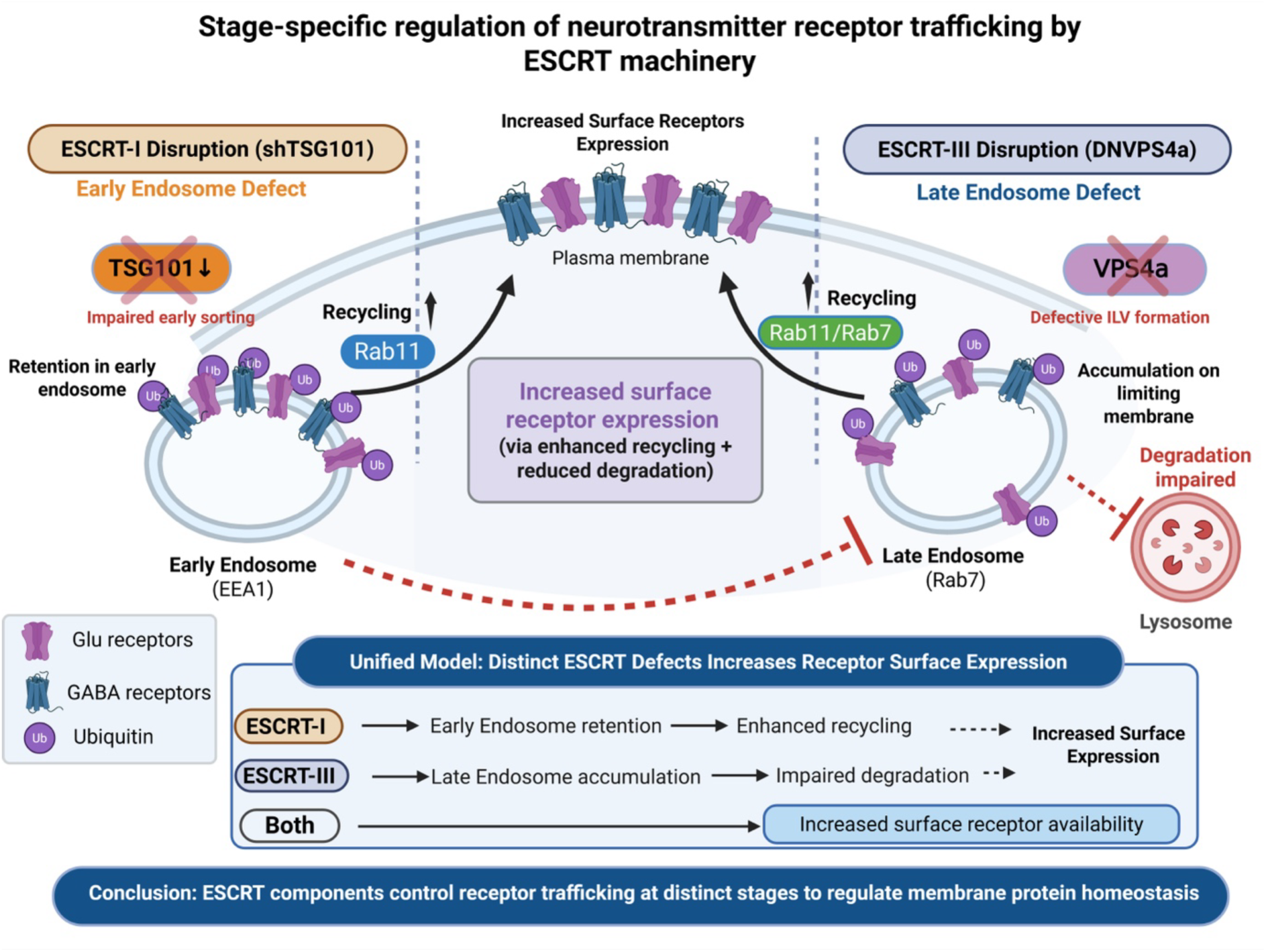

## Introduction

Neurotransmitter receptors at the neuronal surface undergo continuous cycles of endocytosis, intracellular sorting, recycling, and degradation that collectively determine their surface availability and influence synaptic signalling [1]. Rather than functioning as static membrane components, excitatory and inhibitory receptors are dynamically regulated through endosomal trafficking pathways that control receptor localisation, turnover, and responsiveness to neuronal activity [1]. Following internalisation, receptors enter the endosomal system, where they may be recycled back to the plasma membrane or directed toward degradation [2]. This dynamic regulation has been described for multiple receptor classes, including ionotropic glutamate receptors such as α-amino-3-hydroxy-5-methyl-4-isoxazolepropionic acid receptors (AMPA receptors) and N-methyl-D-aspartate receptors (NMDA receptors), which undergo clathrin-mediated endocytosis and subsequent sorting within early and recycling endosomes [3–5]. Similarly, γ-aminobutyric acid type A receptors (GABA_A_) are internalised and trafficked through endosomal compartments in a manner that shapes their steady-state surface levels [3–5]. Across these receptor families, post-endocytic sorting represents a key determinant of receptor distribution, yet the general molecular machinery governing these pathways remains unknown. Dysregulation of these trafficking pathways has been linked to synaptic dysfunction and neurological disease, highlighting the importance of intracellular sorting mechanisms in neuronal homeostasis [6].

Endosomal sorting represents a critical decision point in membrane protein trafficking, directing cargo toward recycling, retention within endosomal compartments, or degradation [7]. The endosomal sorting complexes required for transport (ESCRT) are central components of this system and have been extensively studied for their roles in membrane remodelling and cargo handling in non-neuronal contexts [8–10]. The ESCRT system comprises several functionally distinct complexes that act sequentially during endosomal processing. ESCRT-I components, including tumour susceptibility gene 101 (TSG101), are primarily involved in cargo recognition and sorting of ubiquitinated membrane proteins, whereas ESCRT-III components mediate membrane remodeling and vesicle scission through ATPase-dependent disassembly mechanisms involving vacuolar protein sorting-associated protein 4A (VPS4a) [11, 12]. Although ESCRT proteins are expressed in neurons and have been implicated in dendritic endosome organisation [13, 14], their contribution to neurotransmitter receptor regulation in neurons remains less clearly defined.

Emerging evidence suggests that ESCRT proteins contribute to neuronal membrane organisation, dendritic endosome function, and synaptic protein homeostasis. Alterations in ESCRT pathways have also been associated with neurodegenerative disorders and synaptic dysfunction, supporting a broader role for endosomal trafficking in nervous system physiology and disease [15]. However, several important questions remain unresolved. It is unclear whether ESCRT-dependent trafficking mechanisms regulate neurotransmitter receptor classes uniformly or whether individual receptor subtypes display selective sensitivity to ESCRT perturbation. Moreover, ESCRT complexes comprise multiple components with specialised functions, raising the possibility that ESCRT-I and ESCRT-III may contribute differently to receptor trafficking outcomes [16]. Few studies have addressed whether modulation of specific ESCRT components alters the surface abundance of neurotransmitter receptors, and even fewer have compared excitatory and inhibitory receptor families in this context. The extent to which receptor- or subunit-specific features influence ESCRT-dependent sorting also remains unclear. These gaps limit our understanding of how intracellular sorting machinery shapes the molecular composition of neuronal membranes and highlight the need for targeted analyses of ESCRT function in receptor trafficking pathways.

In this study, we examined how modulation of individual ESCRT components influences neurotransmitter receptor surface expression in heterologous cells and primary neurons. We focused on TSG101, a core ESCRT-I protein involved in cargo recognition, and VPS4a, an ATPase required for ESCRT-III disassembly and endosomal maturation. Using surface biotinylation and imaging-based assays, we assessed how perturbation of these components influences the surface abundance of glutamatergic and GABAergic receptor subtypes in heterologous cells. To evaluate the relevance of these mechanisms in neuronal environments, we extended key analyses to primary neuronal cultures. By comparing the effects of ESCRT-I and ESCRT-III perturbation across receptor classes, we aimed to determine whether ESCRT-dependent regulation is component-specific, receptor-specific, or both. This study centres on receptor surface localisation and post-endocytic sorting, providing a mechanistic framework for how distinct ESCRT components influence receptor trafficking.

## Materials and Methods

### Cell Culture and Transfection

#### HEK293 Cells

Human embryonic kidney 293 (HEK293) cells (ATCC CRL-1573) were maintained in DMEM (DMEM; Gibco) supplemented with 10% fetal bovine serum (Gibco, Thermo Fisher Scientific, USA), 1% penicillin–streptomycin (Gibco, USA), and 2 mM Glutamax (Gibco, USA) at 37 °C in a humidified atmosphere containing 5% CO₂. Cells were passaged every 2–3 days and used between passages 5–20. For receptor trafficking experiments, cells were seeded onto poly-D-lysine-coated plates or coverslips 24 h prior to transfection. Cells were transiently transfected using Lipofectamine 3000 (Thermo Fisher Scientific, USA) according to the manufacturer’s instructions. Total plasmid DNA was kept constant across all experimental conditions by supplementation with empty vector plasmids where necessary.

For glutamate receptor experiments, HEK293 cells were transfected with combinations of NMDA receptor subunits (GluN1/GluN2A), AMPA receptor subunits (GluA1/GluA2), or kainate receptor subunit GluK2 subunits together with scramble shRNA, shRNA targeting TSG101 (shTSG101), wild type TSG101, wild-type VPS4a, or dominant-negative VPS4a (DN-VPS4a; E228Q). For inhibitory receptor experiments, cells were transfected with GABAB receptor subunits GABABR1 and GABABR2 under equivalent ESCRT modulation conditions.

#### Primary Neuronal Cultures

Primary cortical and hippocampal neuronal cultures were prepared from embryonic day 18 Wistar Han rat embryos using standard dissociation procedures ([17]). Briefly, embryonic cortices and hippocampi were dissected in ice-cold Hank’s balanced salt solution and dissociated by gentle trituration. Cells were plated on poly-D-lysine–coated coverslips (Sigma-Aldrich, St. Louis, MO, USA) at 80,000–120,000 cells/well and maintained in Neurobasal medium (Gibco, USA) supplemented with B27 (Gibco, USA), GlutaMAX (Gibco, USA), and penicillin–streptomycin (Gibco, USA) at 37°C in 5% CO₂. Half-medium changes were performed every 3–4 days. Neurons were maintained in culture until DIV10–DIV14 for all experiments.

#### Lentiviral Constructs and Neuronal Transduction

Primary neurons were transduced at DIV7 with lentiviral constructs encoding shTSG101, DN-VPS4a, or respective controls. Lentiviral transfer vectors were obtained from VectorBuilder and included shTSG101, scrambled shRNA controls, recombinant VPS4a constructs, and fluorescent reporter constructs. Viral production followed established second-generation packaging methods ([18]). For transduction, equivalent viral titres were applied directly to neuronal culture medium (1:500–1:1000 dilution). Transduction efficiency was monitored by EGFP or mCherry fluorescence.

Neurons were fixed or lysed for subsequent biochemical and imaging analyses between DIV10 and DIV14.

### Plasmids and shRNA Constructs

A comprehensive set of plasmids were used to manipulate neurotransmitter receptor expression and ESCRT function. All constructs were sequence-verified and maintained according to standard molecular biology procedures. Receptor expression plasmids included GluK2-GFP (JM Henley lab), GluN1A-pEGFP (Addgene #17926), GluN2A (JM Henley lab), GluN2B-pEGFP (Addgene #17925), GluA1 and GluA2 (JM Henley lab). GABA receptor constructs included GABABR1-Myc and GABA_B_R2-HA (JM Henley lab), and Myc-GABA_A_RG2 (Addgene #119730).

ESCRT-related constructs included wild-type TSG101-myc (JM Henley lab), pLNCX2-mCherry-VPS4a (Addgene #115334), and the ATPase-deficient dominant-negative VPS4a mutant E228Q-VPS4a-HA (Addgene #200087). Lentiviral transfer vectors were obtained from VectorBuilder and included rTSG101 (VB240813-1424gea), rVPS4a (VB240809-1242ajb), DN-VPS4a (VB240813-1403ubj), scrambled shRNA (VB200120-1037smy), and shTSG101 (VB900077-7127vsc). Fluorescent reporter controls (VB240813-1423gnb) and empty vector plasmids (pCAT) were used for baseline expression and transduction tracking.

### Lentiviral Production

Lentiviral particles were generated in HEK293T cells using a second-generation packaging system. Transfer vectors included control EGFP/mCherry reporters, recombinant rTSG101 and rVPS4a constructs, dominant-negative VPS4a (E228Q), scrambled shRNA, and shTSG101. HEK293T cells were seeded at 70–80% confluence in 10-cm dishes and co-transfected using Lipofectamine 3000 (Thermo Fisher Scientific, USA) with the following plasmids at a 45:20:15:20 mass ratio: Transfer vector (gene of interest)/ PLP1 (gag/pol)/ PLP2 (rev)/ pVSV-G (envelope glycoprotein).

DNA–lipid complexes were prepared in Opti-MEM (Gibco, USA) and applied to cells for 6 h before replacing with fresh medium. Viral supernatants were collected at 48 h and 72 h, clarified by 0.45 µm filtration (Millipore, Burlington, MA, USA), aliquoted, and stored at −80 °C. Viral titres were estimated by transduction efficiency in HEK293T cells, and equivalent titres were used across all neuronal experiments.

### Surface Biotinylation Assays

Surface receptor expression was quantified using sulfo-NHS-SS-biotin (Thermo Fisher, USA) following established methods for membrane protein isolation ([19]). Cells were washed with ice-cold PBS (Sigma-Aldrich, USA) and incubated with 0.5 mg/mL biotin per HBSS (Gibco, USA) for 30 min at 4 °C. Excess reagent was quenched with 100 mM glycine. Lysates were prepared in RIPA buffer with protease inhibitor cocktail (Roche, Basel, Switzerland), and biotinylated proteins were isolated using Pierce™ High-Capacity Streptavidin Agarose (Thermo Fisher, USA). Surface and total fractions were analysed by SDS-PAGE and immunoblotting. GAPDH was used as a loading control for total lysates and was absent from surface fractions.

### Immunoblotting

Proteins were resolved on 8% SDS-PAGE gels (Bio-Rad, Hercules, CA, USA) and transferred to PVDF membranes (Millipore, USA). Membranes were blocked in 5% BSA (Sigma-Aldrich, St. Louis, MO, USA) and incubated with primary antibodies against GluN1 (mouse, 1A4B10, Synaptic Systems), GluN2A (rabbit, AB1555P, Synaptic Systems), GluA1 (rabbit, 182003, Synaptic Systems), GluA2 (rabbit, 182103, Synaptic Systems), GluK2 (mouse, 1A2B3C, Synaptic Systems), GABRG2 (rabbit, AB_223344, Abcam), GABABR1 (guinea pig, AB_11234, Abcam), TSG101 (rabbit, Ab30871, Abcam), VPS4a (mouse, SAB4200215, Sigma-Aldrich), EGFR (rabbit, AB52894, Abcam), ubiquitin (mouse, P4D1, Sigma-Aldrich; rabbit, U5379, Sigma-Aldrich), and β-actin (mouse, AC-15, Sigma-Aldrich). Fluorescent secondary antibodies (IRDye 680RD anti-mouse IgG, 925-68070; IRDye 800CW anti-rabbit IgG, 925-32211; LI-COR Biosciences, Lincoln, NE, USA) were used for detection on a LI-COR imaging system Odyssey DLx. Band intensities were quantified using ImageJ (Fiji, version 2.16.0/1.54p as described previously ([20]).

### Immunocytochemistry

Surface receptor localisation was assessed using non-permeabilised immunocytochemistry following established protocols for extracellular epitope detection ([21]). Cells were fixed in 4% paraformaldehyde (Sigma-Aldrich, USA) without detergent. Primary antibodies recognising extracellular domains of GluN1, GluN2A, GluA1, GluA2, GluK2, GABA_A_RG2, or GABA_B_R1 were applied, followed by Alexa Fluor–conjugated secondary antibodies (Thermo Fisher Scientific, USA). Neuronal morphology and transduction were visualised via EGFP or mCherry expression.

Images were acquired using a Zeiss LSM confocal microscope (Carl Zeiss AG, Oberkochen, Germany) with a 63× oil objective (NA 1.3). Laser power, gain, and offset were kept constant across conditions. Surface fluorescence intensity and colocalisation were quantified in ImageJ. For neuronal analyses, n = 6- 8 neurons per condition were evaluated from 3 independent biological cultures.

### Statistical Analysis

Data are presented as mean ± SEM. Statistical analyses were performed using GraphPad Prism 10. Comparisons between two groups used unpaired two-tailed Student’s t-tests. Multi-group comparisons used one-way ANOVA with Tukey’s post hoc test. Significance thresholds were defined as p < 0.05, p < 0.01, p < 0.001, and p < 0.0001. For biochemical assays, n = 3-4 independent experiments; for immunocytochemistry, n = 6-8 neurons per condition.

## Results

### Validation of ESCRT Perturbation by TSG101 Knockdown and Dominant-Negative VPS4a Expression

To investigate the contribution of ESCRT machinery to neurotransmitter receptor trafficking, we first validated the experimental modulation of ESCRT-I and ESCRT-III pathways in HEK293 cells. ESCRT-I function was disrupted using shRNA-mediated knockdown of TSG101, whereas ESCRT-III-associated activity was perturbed through expression of an ATPase-deficient dominant-negative VPS4a mutant (DN-VPS4a).

Immunoblot analysis demonstrated that increasing amounts of shTSG101 produced a dose-dependent reduction in endogenous TSG101 expression (**Fig. S1A–D**). Consistent with the established role of ESCRT-I in endosomal cargo sorting, TSG101 depletion was associated with altered expression profiles of epidermal growth factor receptor (EGFR), a canonical ESCRT-dependent cargo protein. In contrast, expression levels of VPS4a and GAPDH remained comparatively stable, indicating that TSG101 knockdown did not produce nonspecific disruption of global protein expression.

Similarly, transfection of increasing amounts of DN-VPS4a resulted in graded accumulation of mutant VPS4a protein without overt effects on total protein loading controls (**Fig. S1E–H**). ESCRT perturbation was further associated with accumulation of ubiquitinated proteins, consistent with impaired endosomal sorting and defective degradative trafficking (**Fig. S2A–D**). Together, these findings confirm effective and controlled disruption of ESCRT function under the experimental conditions used throughout this study.

Based on these validation experiments, intermediate perturbation conditions were selected for subsequent analyses of neurotransmitter receptor trafficking and surface expression.

### TSG101 Knockdown Alters Glutamate Receptor Surface Expression

To determine whether ESCRT-I contributes to glutamatergic receptor trafficking, we examined the effects of TSG101 knockdown on the surface expression of NMDA, AMPA, and kainate receptor subunits in HEK293 cells using surface biotinylation assays.

Reduction of TSG101 significantly increased the surface abundance of NMDA receptor subunits GluN1 and GluN2A relative to scramble shRNA controls (**Fig. 1A–C**). Although modest alterations in total receptor levels were also observed, changes in surface-associated receptor fractions were proportionally greater, indicating selective effects on receptor trafficking and membrane localisation.

**Fig. 1.**
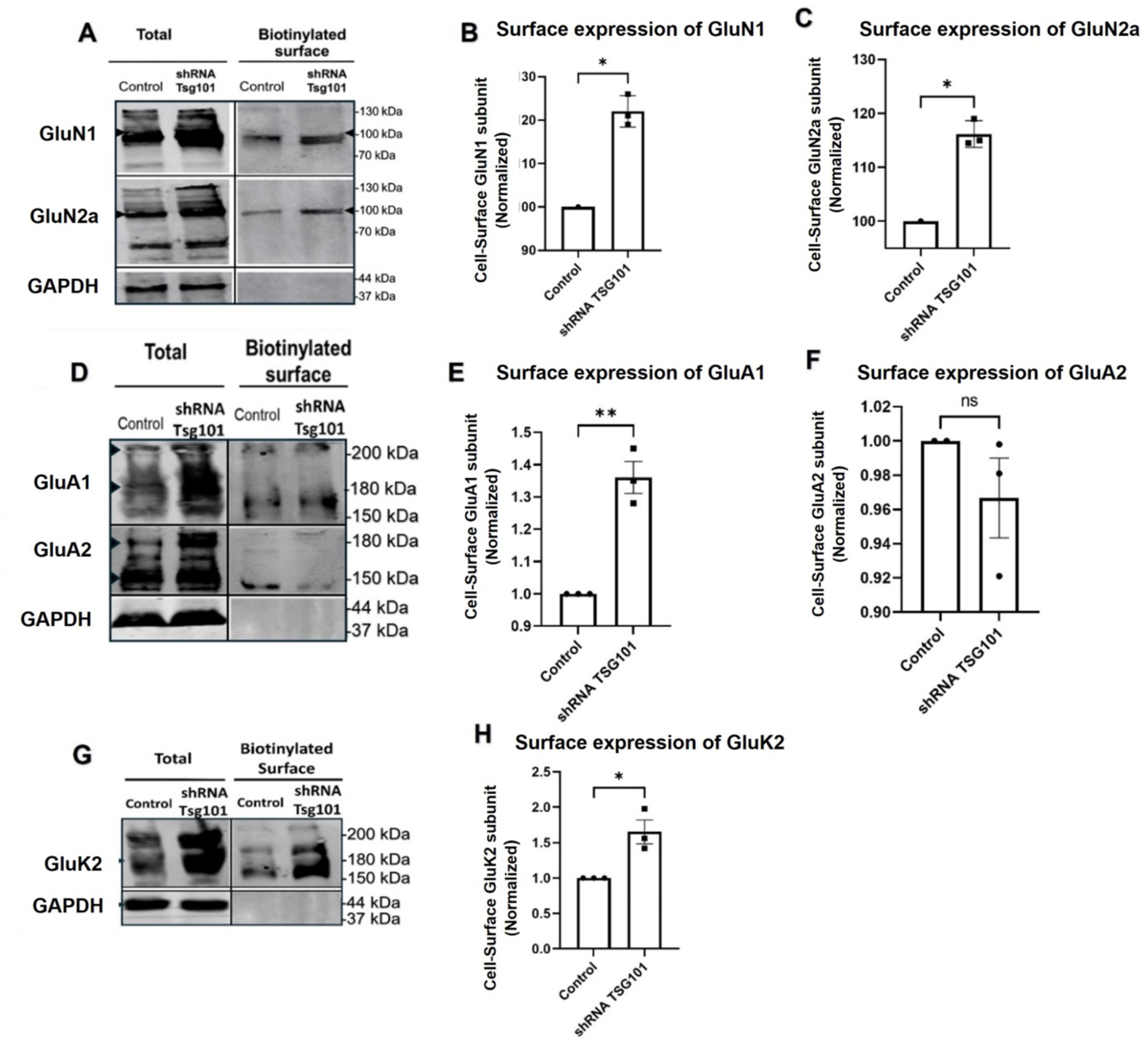
TSG101 knockdown selectively alters glutamate receptor surface expression. (A) Representative immunoblots showing total and surface-biotinylated NMDA receptor subunits GluN1 and GluN2A in HEK293 cells transfected with scramble shRNA (control) or shTSG101. Surface proteins were isolated by sulfo-NHS-SS-biotin-based biotinylation and analysed by immunoblotting. GAPDH served as a loading control for total lysate fractions and was absent from biotinylated fractions. (B–C) Quantification of surface GluN1 (B) and GluN2A (C) normalised to total receptor levels and expressed as fold change relative to scramble shRNA. TSG101 knockdown significantly increased the surface abundance of both NMDA receptor subunits. (D) Representative immunoblots showing total and surface-biotinylated AMPA receptor subunits GluA1 and GluA2. (E–F) Quantification of surface GluA1 (E) and GluA2 (F) normalised to total receptor levels. TSG101 knockdown increased GluA1 surface expression; GluA2 surface levels were not significantly altered. (G) Representative immunoblots showing total and surface-biotinylated GluK2. (H) Quantification of GluK2 surface expression normalised to total protein, showing increased surface abundance following TSG101 knockdown. Data are presented as mean ± SEM (n = 3 independent experiments). Unpaired two-tailed Student’s t-test; *p < 0.05, **p < 0.01; ns, not significant.

AMPA receptor subunits displayed differential sensitivity to ESCRT-I disruption. Surface expression of GluA1 was significantly increased following TSG101 knockdown, whereas GluA2 surface levels showed comparatively smaller or nonsignificant changes under equivalent conditions (**Fig. 1D–F**). These findings indicate subunit-specific regulation of AMPA receptor trafficking by ESCRT-I machinery. Similarly, TSG101 knockdown increased the surface abundance of the kainate receptor subunit GluK2 (**Fig. 1G–H**). Enhanced surface localisation was accompanied by detectable changes in total receptor expression, although the magnitude of surface accumulation exceeded changes in total protein abundance.

Collectively, these findings demonstrate that disruption of ESCRT-I differentially regulates glutamate receptor surface expression in a receptor- and subunit-dependent manner, and changes in surface expression were not uniformly proportional to changes in total protein levels.

### Perturbation of VPS4a Produces Distinct Alterations in Glutamate Receptor Surface Localisation

To determine whether ESCRT-III contributes similarly to receptor trafficking regulation, we next examined the effects of dominant-negative VPS4a expression on glutamate receptor surface localisation using non-permeabilised immunocytochemistry.

Expression of DN-VPS4a increased surface-associated staining of NMDA receptor subunits GluN1 and GluN2A compared with wild-type VPS4a controls (**Fig. 2A–B**). Although the direction of change resembled that observed following TSG101 depletion, the magnitude and spatial distribution of receptor accumulation differed between ESCRT-I and ESCRT-III perturbation conditions.

**Fig. 2.**
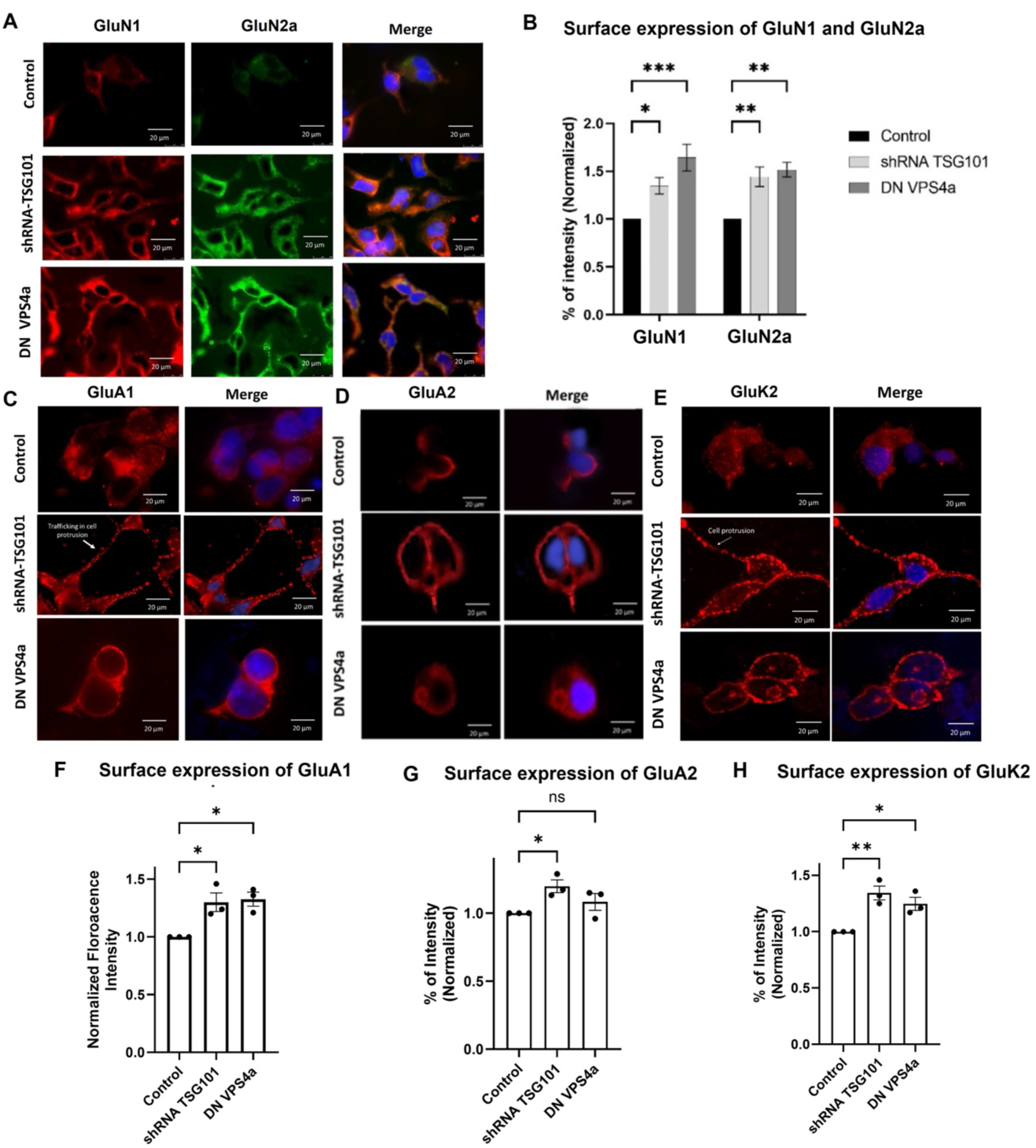
ESCRT modulation alters surface localisation of glutamate receptor subunits in HEK293 cells. (A) Representative immunocytochemical images showing surface staining of GluN1 (red) and GluN2A (green) in HEK293 cells under control conditions, following shTSG101, or expression of dominant-negative VPS4a (DN-VPS4a). Merged images include nuclear staining (blue). Scale bar: 20 µm. (B) Quantification of surface GluN1 and GluN2A fluorescence intensity normalised to control. Both shTSG101 and DN-VPS4a significantly altered surface expression of GluN1 and GluN2A. (C–E) Representative images of surface GluA1 (C), GluA2 (D), and GluK2 (E) staining under control, shTSG101, and DN-VPS4a conditions. Scale bars: 20 µm. (F) Quantification of surface GluA1 fluorescence intensity. Both shTSG101 and DN-VPS4a significantly increased GluA1 surface expression. (G) Quantification of surface GluA2 fluorescence intensity. Surface GluA2 showed modest or non-significant changes depending on condition. (H) Quantification of surface GluK2 fluorescence intensity. Both shTSG101 and DN-VPS4a significantly altered GluK2 surface expression. Data are presented as mean ± SEM (n = 3 independent experiments). One-way ANOVA with Tukey’s post hoc test; *p < 0.05, **p < 0.01, ***p < 0.001; ns, not significant.

AMPA receptor subunits again exhibited differential sensitivity. Surface-associated GluA1 immunoreactivity was increased following DN-VPS4a expression, whereas GluA2 showed comparatively modest alterations (**Fig. 2C–D**). In some cells, altered GluA2 localisation within membrane protrusions and peripheral compartments was observed following VPS4a disruption.

Similarly, DN-VPS4a expression increased surface-associated GluK2 staining relative to controls (**Fig. 2E–F**). Compared with TSG101 knockdown, VPS4a perturbation produced broader intracellular accumulation patterns consistent with altered endosomal maturation and impaired degradative trafficking.

Together, these findings indicate that ESCRT-III disruption alters glutamate receptor surface localisation through mechanisms that partially overlap with, but remain distinct from, those associated with ESCRT-I perturbation.

### TSG101 Knockdown Differentially Regulates GABA_B_ Receptor Surface Expression

To determine whether ESCRT-dependent trafficking regulation extends to inhibitory receptors, we next examined the effects of TSG101 knockdown on GABA_B_ receptor surface expression.

Surface biotinylation assays revealed that TSG101 depletion significantly increased surface-associated GABA_B_R1 expression relative to scramble shRNA controls (**Fig. S3A–C**). In contrast, GABA_B_R2 displayed comparatively modest changes in surface abundance despite detectable alterations in total receptor expression.

Analysis of surface-to-total receptor ratios further demonstrated differential regulation between GABA_B_ receptor subunits, suggesting that ESCRT-I disruption selectively influences receptor trafficking efficiency rather than uniformly altering total receptor abundance.

These findings indicate that ESCRT-I-dependent trafficking mechanisms are not restricted to glutamatergic receptors but also contribute to inhibitory metabotropic receptor surface regulation in a subunit-specific manner.

### ESCRT Perturbation Differentially Regulates Neurotransmitter Receptor Surface Localisation in Primary Neurons

To determine whether the trafficking phenotypes observed in heterologous systems are conserved in neuronal contexts, we next examined endogenous neurotransmitter receptor localisation in primary cortical and hippocampal neurons following ESCRT perturbation.

### NMDA Receptors

TSG101 knockdown significantly increased the surface-to-total ratios of GluN1 and GluN2A in primary neurons, as determined by surface biotinylation assays (**Fig. 3A–D**). Immunocytochemical analyses further demonstrated enhanced dendritic surface localisation of NMDA receptor subunits following shTSG101 expression (**Fig. 3G–H**).

**Fig. 3.**
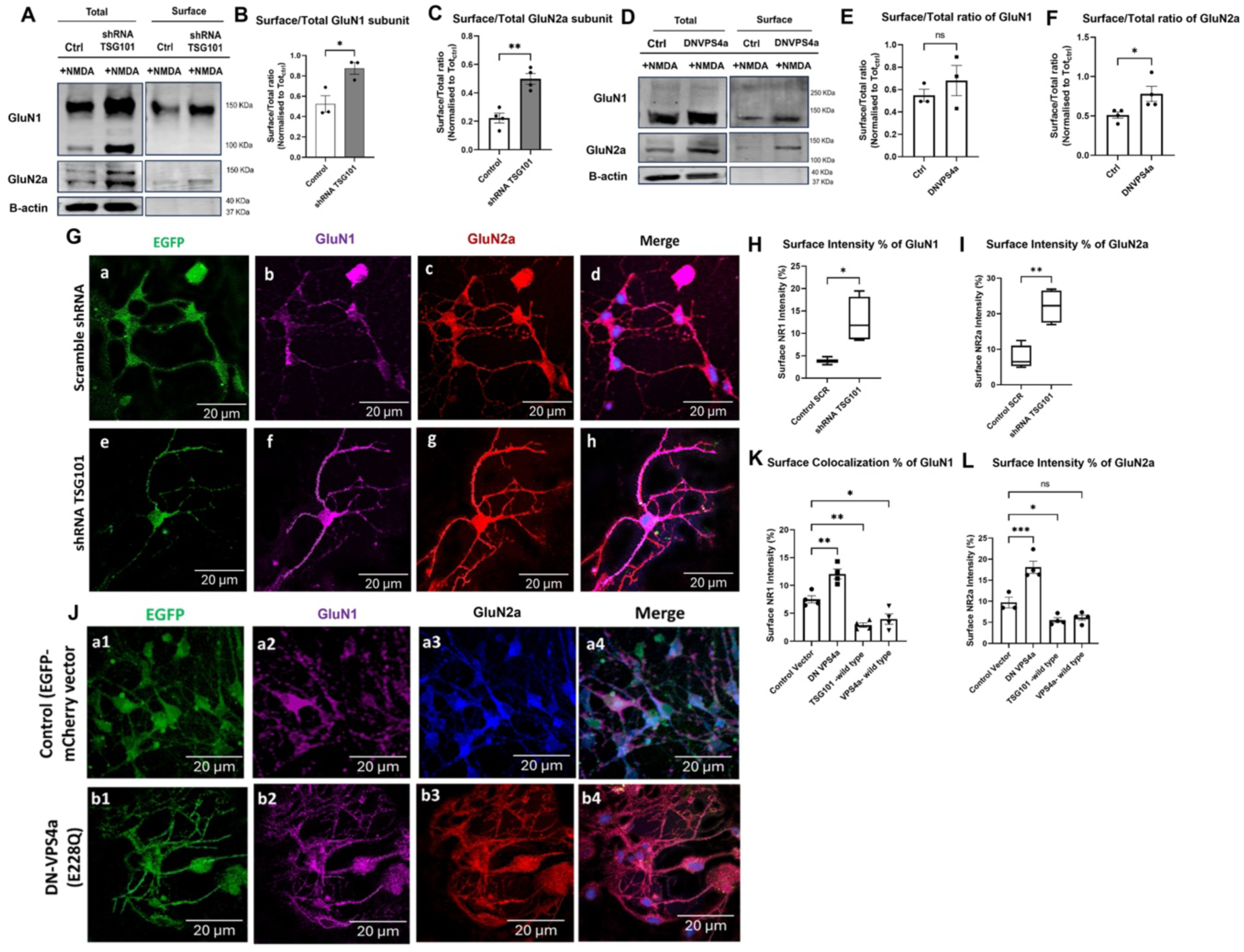
NMDA receptor surface expression increases upon ESCRT pathway disruption in primary cortical neurons. (A–C) Representative immunoblots and quantification showing increased surface/total ratios of GluN1 and GluN2A in primary cortical neurons (DIV14) following shTSG101 transduction. β-actin served as a loading control. (D–F) Representative immunoblots and quantification showing increased GluN1 and GluN2A surface/total ratios following DN-VPS4a expression. (G–H) Representative immunocytochemical images and quantification of surface GluN1 and GluN2A under non-permeabilised conditions in neurons expressing scramble shRNA (control) or shTSG101. EGFP indicates transduction (green). Scale bars: 20 µm. (I–L) Representative immunocytochemical images and quantification of surface GluN1 and GluN2A in neurons expressing control vector or DN-VPS4a. Scale bars: 20 µm. Data are presented as mean ± SEM. For biochemical assays, n = 3 independent experiments; for immunocytochemistry, n = 6–8 neurons per condition from 3 independent cultures. One-way ANOVA with Tukey’s post hoc test; *p < 0.05, **p < 0.01, ***p < 0.001; ns, not significant.

Expression of DN-VPS4a similarly increased surface-associated GluN1 and GluN2A staining; however, receptor distribution patterns differed from those observed following TSG101 depletion (**Fig. 3I–L**), suggesting distinct effects of ESCRT-I and ESCRT-III disruption on neuronal receptor trafficking.

### AMPA Receptors

AMPA receptor subunits displayed distinct sensitivities to ESCRT perturbation. TSG101 knockdown increased surface GluA1 and GluA2 (**Fig. 4A–C**), with GluA1 showing a more pronounced elevation. Developmental analysis at DIV14 revealed sustained GluA1 increases and a stronger GluA2 response at this stage (**Fig. 4E–F**). Dominant-negative VPS4a produced a different pattern: GluA2 surface levels increased, including enhanced localisation within dendritic protrusions (**Fig. 4J–L**), whereas GluA1 changes were comparatively modest (**Fig. 4G–I**).

**Fig. 4.**
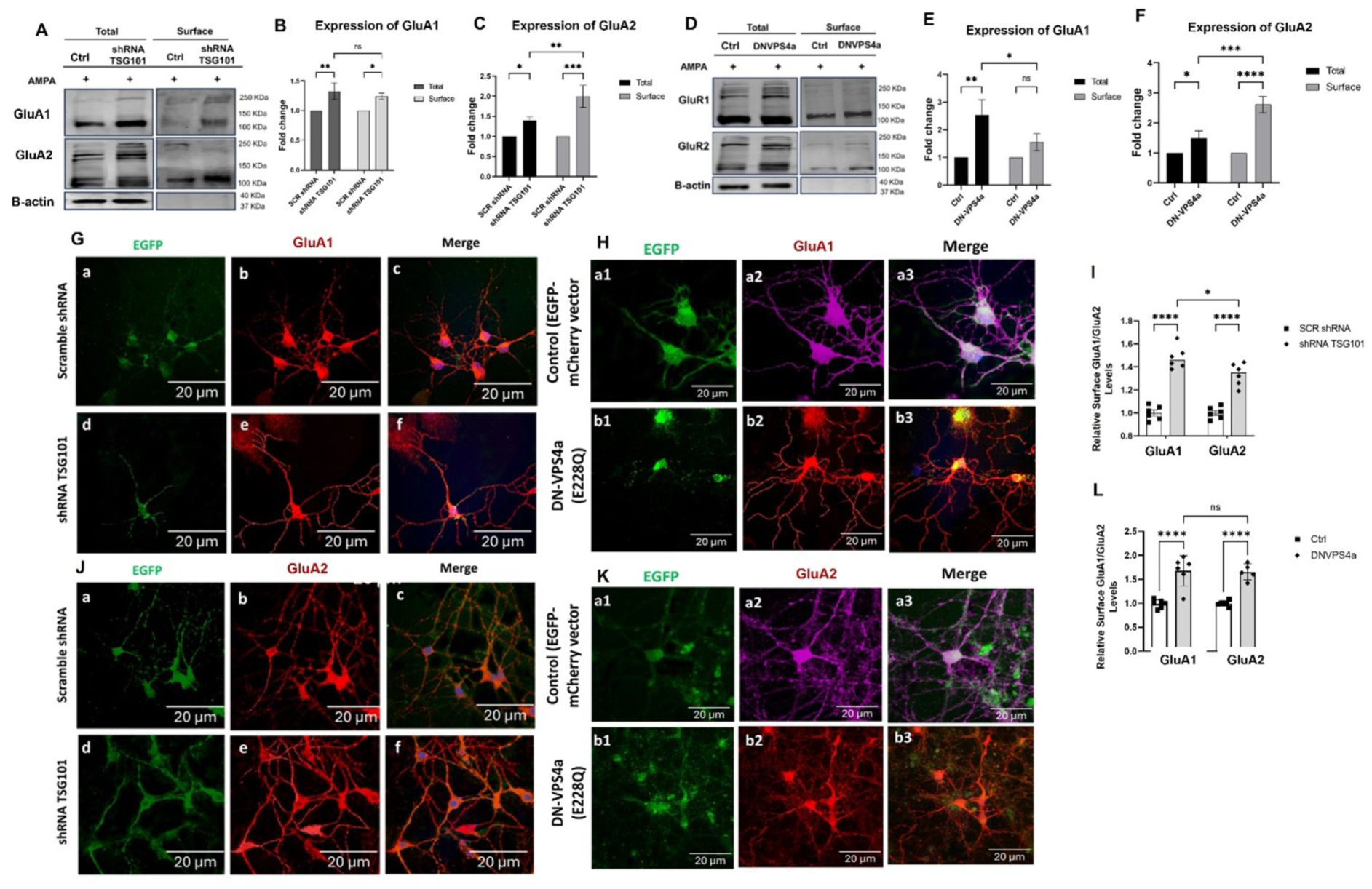
AMPA receptor subunits show differential ESCRT-dependent trafficking in primary cortical neurons. (A–C) Representative immunoblots and quantification showing differential effects of shTSG101 on GluA1 and GluA2 in primary cortical neurons (DIV14). GluA1 displayed robust increases in both total and surface levels, whereas GluA2 showed more modest changes. Neurons were treated with AMPA (20 µmol/L, 20 min) prior to surface biotinylation. β-actin served as a loading control. n = 4 independent cultures. *p < 0.05, **p < 0.01, ***p < 0.001; one-way ANOVA with Tukey’s post hoc test. (D–F) Representative immunoblots and quantification showing preferential effects of DN-VPS4a on GluA1 surface expression compared with GluA2. n = 4. **p < 0.01, ***p < 0.001, ****p < 0.0001; unpaired two-tailed Student’s t-test. (G–H) Representative surface immunocytochemical images of GluA1 in neurons expressing scramble shRNA (G, top) or shTSG101 (G, bottom), and control vector (H, top) or DN-VPS4a (H, bottom). EGFP indicates transduction (green). Scale bars: 20 µm. (I) Quantification of surface GluA1 and GluA2 intensity following shTSG101 or scramble shRNA. n ≥ 8 neurons per condition. ****p < 0.0001, *p < 0.05; two-way ANOVA. (J–K) Representative surface immunocytochemical images of GluA2 following shTSG101 (J) and DN-VPS4a (K). Scale bars: 20 µm. (L) Quantification of surface GluA1 and GluA2 following DN-VPS4a. n ≥ 8 neurons per condition. ***p < 0.001; ns, not significant; two-way ANOVA. All data presented as mean ± SEM.

### Kainate Receptors

GluK2 surface expression was markedly elevated following TSG101 knockdown, as shown by biotinylation and immunofluorescence (**Fig. 5**). This increase persisted in the presence of kainate stimulation, indicating that the effect was not ligand-dependent. Dominant-negative VPS4a also increased surface GluK2, though to a different extent (**Fig. 5C–D, 5G–H**). These results show that both ESCRT-I and ESCRT-III contribute to the control of kainate receptor surface distribution.

**Fig. 5.**
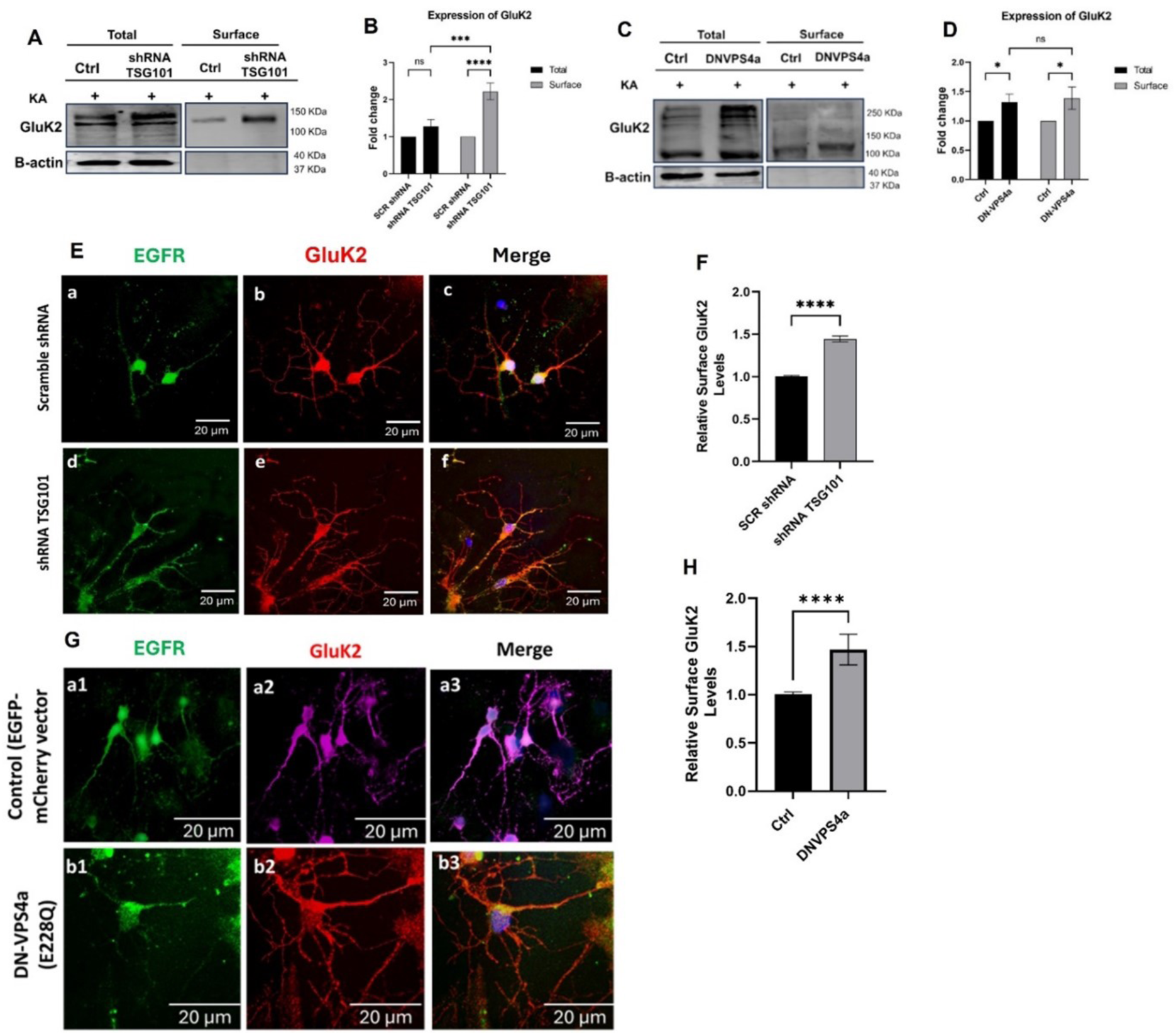
Kainate receptor GluK2 trafficking is ESCRT-dependent in primary cortical neurons. (A–B) Representative immunoblots and quantification showing increased total and surface GluK2 levels in primary cortical neurons (DIV14) following shTSG101 transduction. Neurons were treated with kainate (20 µmol/L, 10 min) prior to surface biotinylation. β-actin served as a loading control. n = 3 independent cultures. ***p < 0.001, ****p < 0.0001; ns, not significant; one-way ANOVA with Tukey’s post hoc test. (C–D) Representative immunoblots and quantification showing increased GluK2 levels following DN-VPS4a expression. n = 4. *p < 0.05; ns, not significant; one-way ANOVA. (E–F) Representative surface immunocytochemical images and quantification of GluK2 in neurons expressing scramble shRNA (E, top) or shTSG101 (E, bottom). EGFP indicates transduction (green). Quantification (F) demonstrates approximately 1.5-fold increase in surface GluK2 with shTSG101. n ≥ 8 neurons per condition. ****p < 0.0001; unpaired two-tailed Student’s t-test. (G–H) Representative surface immunocytochemical images and quantification of GluK2 in neurons expressing control vector (G, top) or DN-VPS4a (G, bottom). Quantification (H) demonstrates approximately 1.4-fold increase in surface GluK2 with DN-VPS4a. n ≥ 8 neurons per condition. ****p < 0.0001; unpaired two-tailed Student’s t-test. Scale bars: 20 µm. All data presented as mean ± SEM.

### GABA Receptors

TSG101 knockdown increased surface levels of the ionotropic GABA_A_ receptor subunit gamma 2 (GABA_A_RG2) and the metabotropic GABA_B_ receptor subunit 1 GABA_B_R1, with biotinylation assays revealing elevated surface-to-total ratios for both subunits (**Fig. 6A–C**). Immunofluorescence analysis showed higher dendritic surface-associated signal for GABA_A_RG2 and GABA_B_BR1 in shTSG101-treated neurons (**Fig. 6G–J**). In neurons expressing dominant-negative VPS4a, surface GABA_A_RG2 levels were increased (**Fig. 6**), whereas GABA_B_R1 showed minimal change relative to controls.

**Fig. 6.**
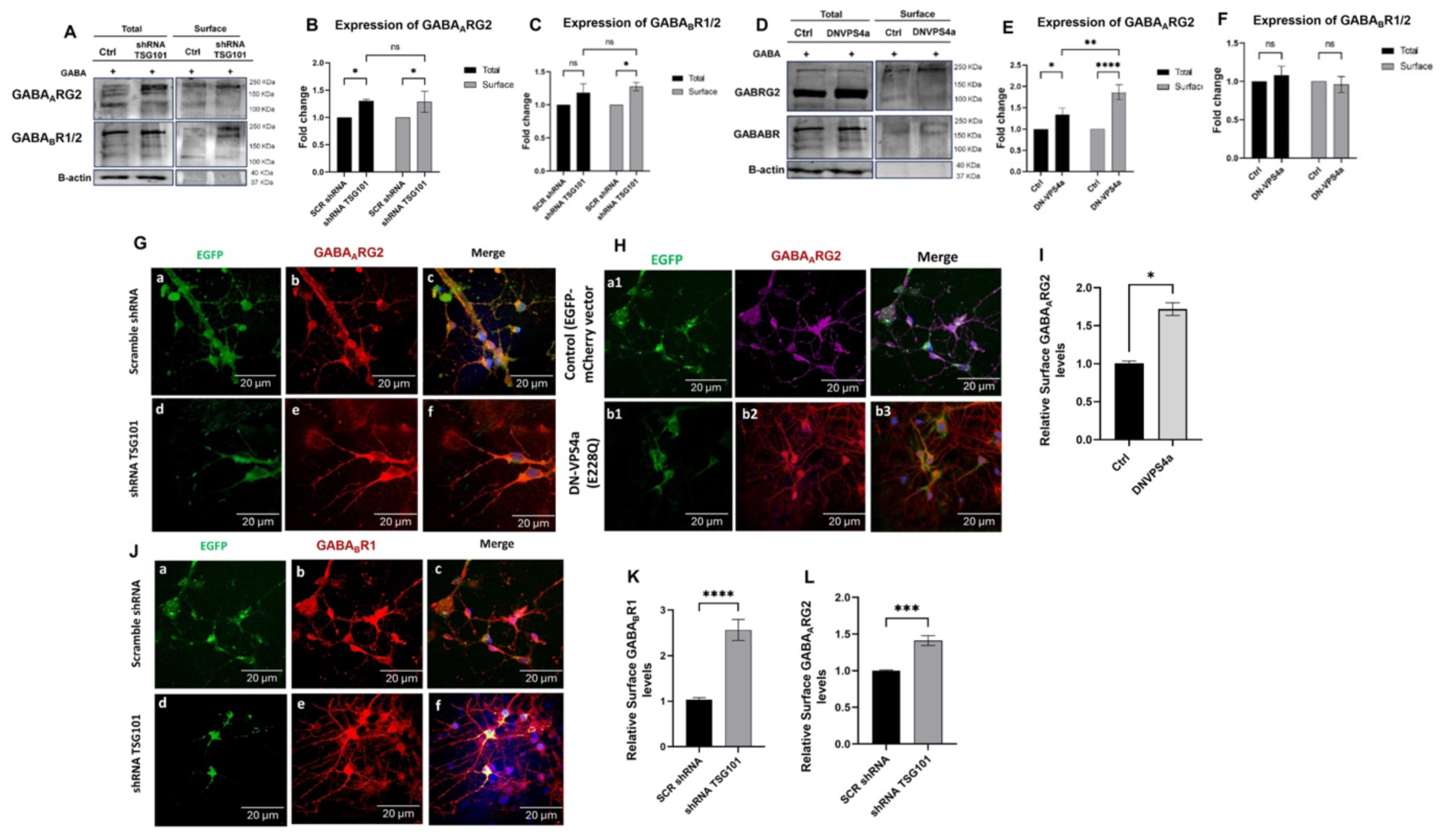
ESCRT-dependent regulation of GABA_A_ and GABA_B_ receptor subunits in primary cortical neurons. (A) Representative surface biotinylation immunoblots showing total and surface levels of GABA_A_RG2 and GABA_B_R1 in primary cortical neurons expressing scramble shRNA or shTSG101. β-actin served as a loading control. (B) Quantification of GABA_A_RG2 expression following shTSG101. Surface GABA_A_RG2 was significantly increased compared with scramble controls; total levels were unchanged. n = 3 independent biological replicates. *p < 0.05; ns, not significant; unpaired two-tailed Student’s t-test. (C) Quantification of GABA_B_R1 expression following shTSG101. Surface GABA_B_R1 was significantly elevated; total protein was unchanged. n = 3 biological replicates. *p < 0.05; ns, not significant; unpaired two-tailed Student’s t-test. (D) Representative surface biotinylation immunoblots showing total and surface GABA_A_RG2 and GABA_B_R1 in neurons expressing DN-VPS4a. β-actin served as a loading control. (E) Quantification of GABA_A_RG2 following DN-VPS4a expression. Surface GABRG2 was significantly increased. n = 3 biological replicates. **p < 0.01, ****p < 0.0001; one-way ANOVA with Tukey’s post hoc test. (F) Quantification of GABA_B_R1 following DN-VPS4a. No significant change in total or surface levels. n = 3 biological replicates. ns, not significant; unpaired two-tailed Student’s t-test. (G–H) Representative immunofluorescence images of neurons stained for GABA_A_RG2 (red) under scramble shRNA (G, top), shTSG101 (G, bottom), and DN-VPS4a (H) conditions. EGFP indicates transduction (green). Scale bars: 20 µm. (I) Quantification of surface GABRG2 in control versus DN-VPS4a neurons. n ≥ 8 neurons from 3 biological replicates. *p < 0.05; unpaired two-tailed Student’s t-test. (J) Representative immunofluorescence images of neurons stained for GABA_B_R1 (red) under scramble shRNA (top) and shTSG101 (bottom) conditions. Scale bars: 20 µm. (K) Quantification of surface GABA_B_R1 in shTSG101 neurons. n ≥ 8 neurons from 3 biological replicates. ****p < 0.0001; unpaired two-tailed Student’s t-test. (L) Quantification of surface GABA_A_RG2 in scramble versus shTSG101 neurons. n ≥ 8 neurons from 3 biological replicates. ***p < 0.001; unpaired two-tailed Student’s t-test. All data presented as mean ± SEM.

### Comparative Analysis Reveals Receptor- and ESCRT-Component-Specific Patterns of Regulation

Across all receptor classes examined, ESCRT perturbation increased receptor surface availability but did not produce uniform trafficking phenotypes. Instead, individual receptor subunits exhibited differential sensitivity to ESCRT-I and ESCRT-III disruption.

To directly compare these patterns across receptor types and ESCRT components, we quantified changes in both surface and total receptor expression following TSG101 knockdown or dominant-negative VPS4a expression (**Figure S4**). This comparative analysis revealed striking receptor- and ESCRT-component-specific patterns. NMDA receptor subunits (GluN1, GluN2A) consistently displayed increased surface localisation following both TSG101 knockdown and VPS4a disruption, although the relative magnitude of these changes differed between ESCRT components. On the other hand, AMPA and GABA receptor subunits exhibited greater variability in response magnitude and spatial distribution. GluA1 and GluK2 were particularly sensitive to ESCRT perturbation, while GluA2 and GABABR2 displayed comparatively selective responses.

Importantly, ESCRT-I and ESCRT-III perturbations produced overlapping yet distinct trafficking phenotypes across receptor classes. TSG101 depletion predominantly increased receptor surface localisation, whereas VPS4a disruption produced broader alterations in receptor distribution and intracellular accumulation patterns. These findings suggest that distinct ESCRT components differentially regulate neurotransmitter receptor trafficking pathways.

### Redistribution of NMDARs Across Endosomal Compartments Following ESCRT Disruption

To investigate how ESCRT perturbation influences intracellular NMDA receptor trafficking, we quantified the colocalisation of the GluN1 subunit with markers of early (EEA1), recycling (Rab11), and late (Rab7) endosomal compartments in dendrites of primary neurons.

Under control conditions, GluN1 displayed detectable colocalisation with EEA1-positive early endosomes, consistent with constitutive receptor internalisation and endosomal sorting (**Fig. 7A-C**). TSG101 knockdown significantly increased GluN1 association with EEA1-positive compartments, as demonstrated by elevated Pearson’s correlation and Manders’ tM2 values (**Fig. 7E–F**). In contrast, neurons expressing dominant-negative VPS4a exhibited reduced GluN1–EEA1 overlap, supported by line-scan analyses (**Fig. 7D**) and significantly decreased Pearson’s correlation and Manders’ tM2 measurements (**Fig. 7E–F**).

**Fig. 7.**
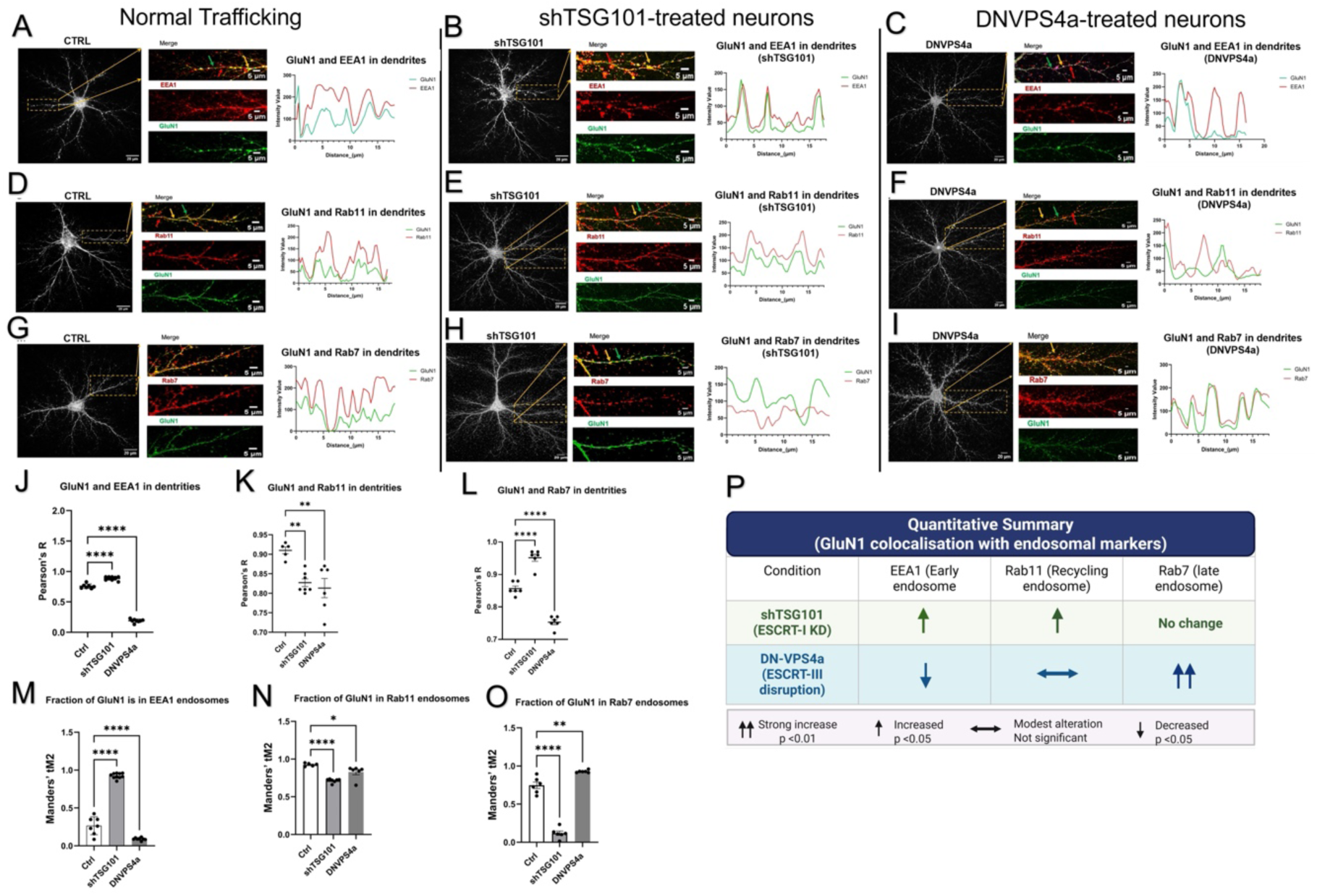
GluN1 redistributes across early, recycling, and late endosomes following ESCRT disruption in primary hippocampal neurons. (A–C) Representative dendritic images showing GluN1 (green) and the early endosome marker EEA1 (red) in control neurons (A), shTSG101-expressing neurons (B), and DN-VPS4a-expressing neurons (C). Merged images with single-channel views and line-scan intensity profiles of GluN1 and EEA1 along dendrites are shown below each condition. Scale bars: 20 µm (overview), 5 µm (insets). (D–F) Representative dendritic images showing GluN1 (green) and the recycling endosome marker Rab11 (red) in control neurons (D), shTSG101-expressing neurons (E), and DN-VPS4a-expressing neurons (F). Line-scan intensity profiles are shown below each condition. Scale bars: 20 µm (overview), 5 µm (insets). (G–I) Representative dendritic images showing GluN1 (green) and the late endosome marker Rab7 (red) in control neurons (G), shTSG101-expressing neurons (H), and DN-VPS4a-expressing neurons (I). Line-scan intensity profiles are shown below each condition. Scale bars: 20 µm (overview), 5 µm (insets). (J–L) Quantification of Pearson’s correlation coefficient (R) for GluN1 colocalisation with EEA1 (J), Rab11 (K), and Rab7 (L) under control, shTSG101, and DN-VPS4a conditions. (M–O) Quantification of Manders’ tM2 values representing the fraction of GluN1 signal within EEA1-positive (M), Rab11-positive (N), and Rab7-positive (O) endosomes. (P) Quantitative summary table of GluN1 colocalisation changes with early (EEA1), recycling (Rab11), and late (Rab7) endosomal markers following shTSG101 or DN-VPS4a. Data represent 3 biological replicates, n = 6 dendrites per condition. All data are presented as mean ± SEM. One-way ANOVA with Tukey’s post hoc test; *p < 0.05, **p < 0.01, ****p < 0.0001; ns, not significant.

GluN1 association with Rab11-positive recycling endosomes was also detected under basal conditions. ESCRT perturbation produced modest but significant alterations in GluN1–Rab11 colocalisation, accompanied by corresponding shifts in line-scan profiles (**Fig. 7J**). By comparison, GluN1 localisation within Rab7-positive late endosomal compartments was markedly increased following dominant-negative VPS4a expression (**Fig. 7K–O**). Line-scan analyses demonstrated pronounced co-alignment of GluN1 and Rab7 fluorescence signals, while both Pearson’s correlation and Manders’ tM2 values were significantly elevated relative to control neurons.

Together, these findings demonstrate that ESCRT-I and ESCRT-III disruption differentially alter GluN1 distribution across the endosomal system, with TSG101 depletion increasing GluN1 association with early endosomal compartments and VPS4a disruption increasing GluN1 localisation within Rab7-positive late endosomes. A schematic summary of GluN1 redistribution across endosomal compartments following ESCRT perturbation is shown in **Fig. 7S**.

## Discussion

The present study demonstrates how distinct ESCRT components regulate neurotransmitter receptor trafficking and the stage-specific endosomal mechanisms that shape receptor surface expression. Using complementary biochemical and imaging-based approaches in heterologous cells and primary neurons, we demonstrate that perturbation of ESCRT-I through TSG101 depletion and disruption of ESCRT-III-associated function through dominant-negative VPS4a expression both increase receptor surface availability, but through distinct intracellular trafficking phenotypes. These findings extend previous studies identifying ESCRT machinery as a key regulator of membrane protein sorting and endosomal organisation [17, 18] and demonstrate that ESCRT-dependent regulation of neurotransmitter receptors is receptor-, subunit-, and pathway-specific.

A major finding of this study is that ESCRT-dependent regulation operates primarily at the level of post-endocytic sorting. Across receptor classes, changes in surface expression were not consistently proportional to alterations in total receptor abundance, indicating that ESCRT perturbation predominantly affects intracellular trafficking decisions rather than receptor synthesis or bulk protein degradation. Although the magnitude of response varied between receptor subunits, this variability is consistent with cargo-selective sorting mechanisms previously described for ESCRT-dependent trafficking pathways [16, 19, 20]. ESCRT-dependent cargo recognition is known to involve ubiquitin-dependent sorting and adaptor-mediated interactions [16], providing a potential basis for the receptor-selective effects observed in the present study. In agreement with this interpretation, ESCRT perturbation was also associated with altered ubiquitin accumulation patterns, further supporting disruption of endosomal cargo processing.

Our findings further demonstrate that ESCRT-I and ESCRT-III perturbations produce non-equivalent trafficking phenotypes that reflect their distinct functional positions within the endosomal pathway. TSG101 depletion increased receptor association with EEA1-positive early endosomal compartments, whereas VPS4a disruption reduced early endosomal association and increased receptor localisation within Rab7-positive late endosomes. These observations are consistent with the established sequential organisation of ESCRT machinery, in which ESCRT-I participates in cargo recognition and early endosomal sorting, whereas ESCRT-III and VPS4 ATPases contribute to membrane remodelling, intraluminal vesicle formation, and endosomal maturation [20–22]. Importantly, the present findings extend these principles to neurotransmitter receptor trafficking and demonstrate that disruption of distinct ESCRT stages alters receptor redistribution across the endosomal system in different ways.

Mechanistically, the increase in receptor surface expression observed following both TSG101 depletion and VPS4a disruption is consistent with altered endosomal trafficking and impaired progression toward degradative pathways. A schematic model summarising these stage-specific trafficking phenotypes is presented in **Fig. 8**.

**Fig. 8.**
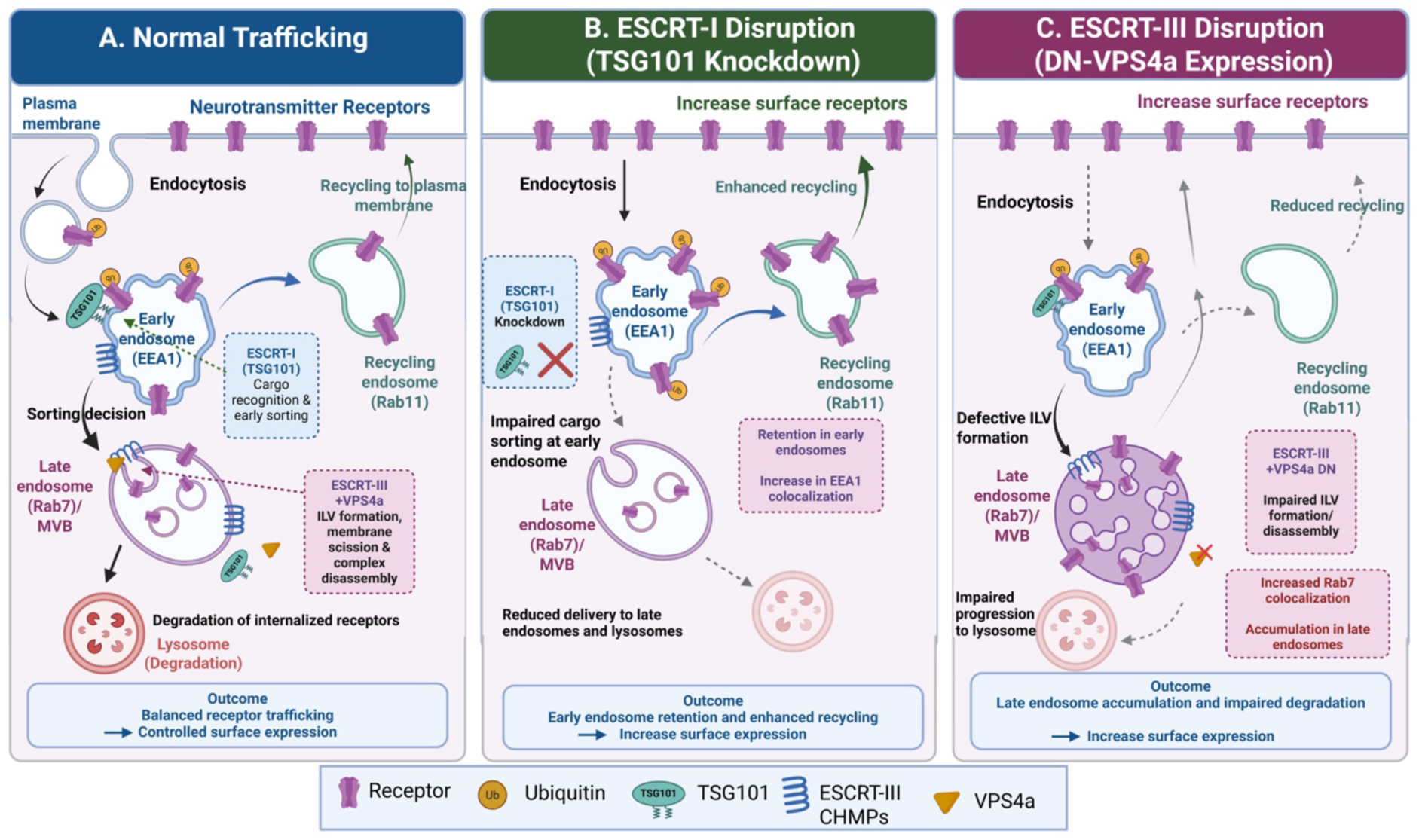
Stage-specific ESCRT regulation of neurotransmitter receptor trafficking. Schematic model illustrating how disruption of distinct ESCRT components differentially alters neurotransmitter receptor trafficking within the endosomal system. (A) Under normal conditions, internalised receptors are sorted within early endosomes and either recycled to the plasma membrane or directed toward late endosomal and lysosomal degradation pathways. (B) ESCRT-I disruption (shTSG101) impairs cargo sorting at early endosomes, resulting in receptor retention within early endosomal compartments and enhanced recycling, thereby increasing steady-state surface receptor expression. (C) ESCRT-III disruption (DN-VPS4a) impairs intraluminal vesicle formation and endosomal maturation, promoting receptor accumulation within Rab7-positive late endosomal compartments and reducing progression through degradative pathways. Despite mechanistically distinct intracellular trafficking defects, both perturbations increase steady-state receptor surface expression. Figure created using BioRender.

Although direct recycling and degradation assays were not performed in the present study, the observed redistribution patterns support a model in which ESCRT dysfunction alters receptor trafficking decisions following endocytosis. Under conditions of ESCRT-I disruption, increased receptor association with early endosomes may reflect altered sorting dynamics that favour receptor retention within recycling-accessible compartments. In contrast, VPS4a disruption produced receptor accumulation within late endosomal compartments, consistent with impaired endosomal maturation and defective cargo progression through degradative pathways [21, 22]. These findings indicate that similar steady-state increases in receptor surface abundance can arise through distinct intracellular trafficking defects within the same endosomal pathway.

Glutamatergic receptor subtypes displayed differential sensitivity to ESCRT perturbation. NMDA receptor subunits consistently exhibited increased surface localisation following both TSG101 knockdown and VPS4a disruption, whereas AMPA receptor subunits displayed greater variability, with GluA1 showing more pronounced changes than GluA2. Similarly, the kainate receptor subunit GluK2 demonstrated robust sensitivity to both ESCRT perturbation paradigms. Previous studies have established that ionotropic glutamate receptors undergo continuous endosomal trafficking that regulates receptor availability, synaptic signalling, and plasticity [23–26]. ESCRT-associated pathways have also been implicated in membrane receptor turnover and synaptic protein homeostasis [27, 28]. The receptor- and subunit-specific effects observed here therefore suggest that distinct glutamate receptor populations may engage different endosomal sorting mechanisms or display variable sensitivity to disruptions in endosomal processing.

Inhibitory receptor subtypes also displayed sensitivity to ESCRT perturbation, although the magnitude and direction of these effects varied between receptor classes and experimental systems. Surface expression of GABA_B_R1 and GABA_A_RG2 was altered following modulation of TSG101 and VPS4a, supporting a broader role for ESCRT-dependent trafficking pathways in the regulation of excitatory and inhibitory receptor balance. Previous studies have demonstrated that inhibitory receptor trafficking is tightly regulated by endosomal sorting and recycling pathways that influence receptor stability and synaptic inhibition [29–31]. The present findings therefore indicate that ESCRT-dependent membrane trafficking mechanisms contribute to the organisation of both excitatory and inhibitory receptor systems.

The conservation of these trafficking phenotypes in primary neurons indicates that ESCRT-dependent regulation of receptor trafficking is not restricted to heterologous systems but represents a broader feature of neuronal membrane organisation. ESCRT proteins have previously been implicated in neuronal endosomal organisation, dendritic maintenance, and synaptic protein homeostasis [27, 32]. At the same time, neuronal analyses revealed additional spatial complexity, including dendritic compartmentalisation and subunit-specific receptor distribution patterns, suggesting that ESCRT-dependent sorting interfaces with neuron-specific trafficking architecture.

Endosomal dysfunction and altered receptor trafficking are increasingly recognized as important contributors to neurological and neuropsychiatric disease [33–35]. ESCRT-associated proteins have been implicated in several neurodegenerative disorders, including frontotemporal dementia, amyotrophic lateral sclerosis, and Alzheimer’s disease, where abnormalities in endosomal organisation and membrane trafficking are common pathological features [36–39]. Altered glutamatergic and GABAergic signalling has also been strongly associated with disorders such as schizophrenia and synaptic dysfunction-related pathologies [40–42]. Although the present study did not directly investigate disease mechanisms, the observed effects of ESCRT perturbation on neurotransmitter receptor localisation support the possibility that impaired endosomal trafficking may contribute to abnormal receptor distribution in disease-associated neuronal dysfunction.

An important strength of this study is the integration of heterologous expression systems with primary neuronal validation. While HEK293 cells enabled controlled examination of receptor-specific trafficking responses, key observations were reproduced in primary neurons, supporting the physiological relevance of these mechanisms in neuronal systems. Nevertheless, several limitations should be acknowledged. The present study focused primarily on receptor localisation and surface expression and did not directly assess the functional consequences of altered receptor trafficking. Electrophysiological recordings, calcium imaging, and receptor activity assays will therefore be important for determining how ESCRT-dependent trafficking influences neuronal signalling and synaptic function. In addition, although redistribution across endosomal compartments was quantified, direct measurements of receptor recycling dynamics and degradative flux were not performed. Future studies examining receptor ubiquitination, adaptor interactions, and live-cell trafficking dynamics will help further define the molecular determinants underlying receptor-selective sensitivity to ESCRT perturbation.

In summary, our data support a unified model in which ESCRT components regulate receptor trafficking through stage-specific control of the endosomal system:

ESCRT-I disruption (TSG101 loss) → early endosomal retention → enhanced recycling

ESCRT-III disruption (VPS4a impairment) → late endosomal accumulation → impaired degradation

Both conditions → increased surface receptor expression via distinct intracellular mechanisms

In conclusion, the present study establishes ESCRT machinery as an important regulator of neurotransmitter receptor surface organisation. ESCRT-I and ESCRT-III perturbations produced distinct effects on endosomal receptor redistribution, resulting in altered surface availability through mechanistically different trafficking phenotypes. These findings highlight the complexity of intracellular membrane sorting pathways in neurons and provide further insight into how ESCRT-dependent trafficking contributes to neurotransmitter receptor homeostasis and neuronal membrane organisation.

## Authors’ Contributions

M.F.S. conceived and designed the experiments, performed all experimental work, analysed and interpreted the data, and drafted the manuscript. S.L.M. contributed to experimental design, data interpretation, and critical revision of the manuscript. S.K. conceived and supervised the study, contributed to experimental design and data interpretation, and revised the manuscript critically for important intellectual content. All authors reviewed and approved the final version of the manuscript.

## Data availability

The datasets generated and/or analysed during the current study are available from the corresponding author upon reasonable request.

## Ethics Statement

All procedures involving animals were approved by the University Animal Welfare and Ethical Review Body (AWERB) and complied with The Animals (Scientific Procedures) Act 1986 (UK).

