## Supplemental figures for "Distinct Endosomal Sorting Complexes Required for Transport Components Differentially Regulate Glutamate and Gamma-Aminobutyric Acid Receptor Surface Expression"

**Running title**: ESCRT Differentially Regulates Receptor Expression

### Supplementary Information

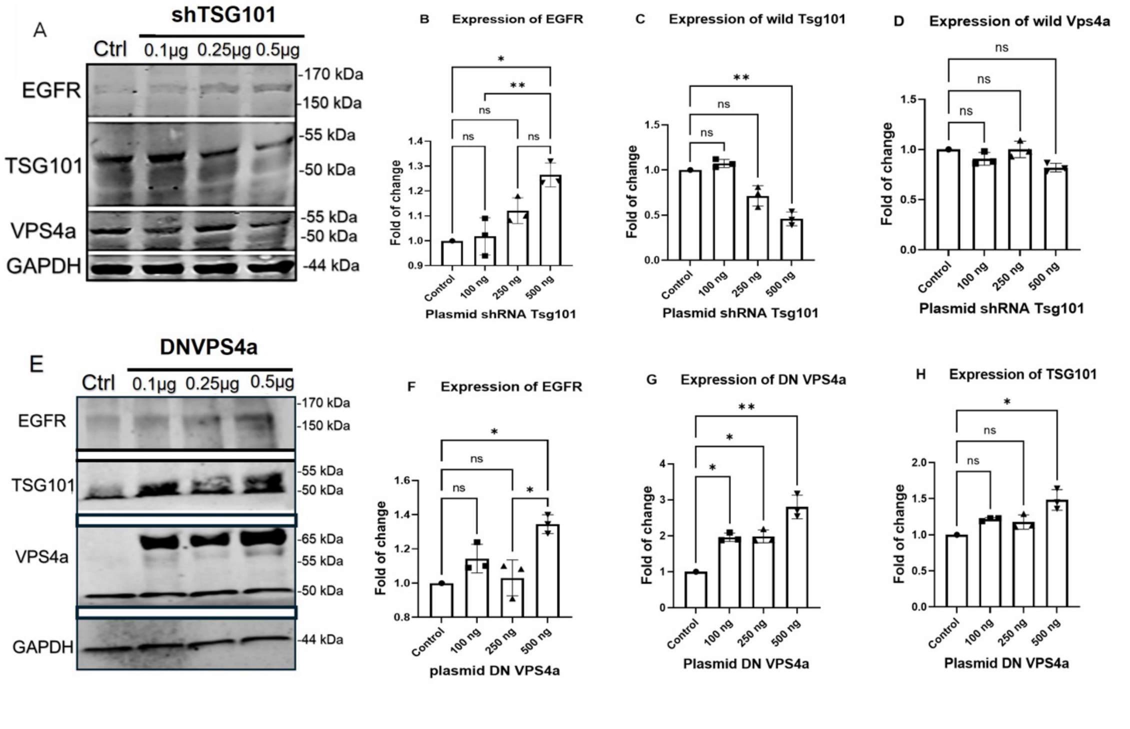


**Fig. S1** Dose-dependent modulation of ESCRT components and effects on EGFR expression in HEK293 cells.

(A) Representative immunoblots showing expression of EGFR, TSG101, VPS4a and GAPDH following transfection with increasing amounts of shRNA targeting TSG101 (0.1, 0.25, and 0.5 µg) compared with control. TSG101 protein levels decrease in a dose-dependent manner, while EGFR and VPS4a expression profiles are shown for comparison. GAPDH was used as a loading control.

(B) Quantification of EGFR protein levels following increasing shRNA TSG101 plasmid amounts, normalised to GAPDH and expressed as fold change relative to control. Data are presented as mean ± SEM ($n = 3$ independent experiments). One-way ANOVA with appropriate post hoc comparisons; *$p < 0.05$, **$p < 0.01$; ns, not significant.

(C) Quantification of endogenous TSG101 levels following shRNA TSG101 transfection, normalised to GAPDH and expressed as fold change relative to control, demonstrating dose-dependent knockdown. Data are presented as mean ± SEM ($n = 3$ independent experiments). One-way ANOVA; **$p < 0.01$; ns, not significant.

(D) Quantification of endogenous VPS4a levels under shRNA TSG101 conditions, normalised to GAPDH and expressed as fold change relative to control. No significant alterations were observed across the tested conditions. Data are presented as mean ± SEM ($n = 3$ independent experiments). One-way ANOVA; ns, not significant.

(E) Representative immunoblots showing expression of EGFR, TSG101, VPS4a and GAPDH following transfection with increasing amounts of dominant-negative (DN) VPS4a plasmid (0.1, 0.25, and 0.5 µg) compared with control. DN-VPS4a expression increases in a dose-dependent manner, with associated changes in EGFR and TSG101 expression profiles shown.

(F) Quantification of EGFR protein levels following DN-VPS4a expression, normalised to GAPDH and expressed as fold change relative to control. Data are presented as mean ± SEM ($n = 3$ independent experiments). One-way ANOVA with post hoc testing; *$p < 0.05$; ns, not significant.

(G) Quantification of DN-VPS4a expression levels following increasing plasmid amounts, normalised to GAPDH and expressed as fold change relative to control, confirming dose-dependent expression. Data are presented as mean ± SEM ($n = 3$ independent experiments). One-way ANOVA; *$p < 0.05$, **$p < 0.01$.

(H) Quantification of endogenous TSG101 levels following DN-VPS4a expression, normalised to GAPDH and expressed as fold change relative to control. Data are presented as mean ± SEM ($n = 3$ independent experiments). One-way ANOVA; *$p < 0.05$; ns, not significant.

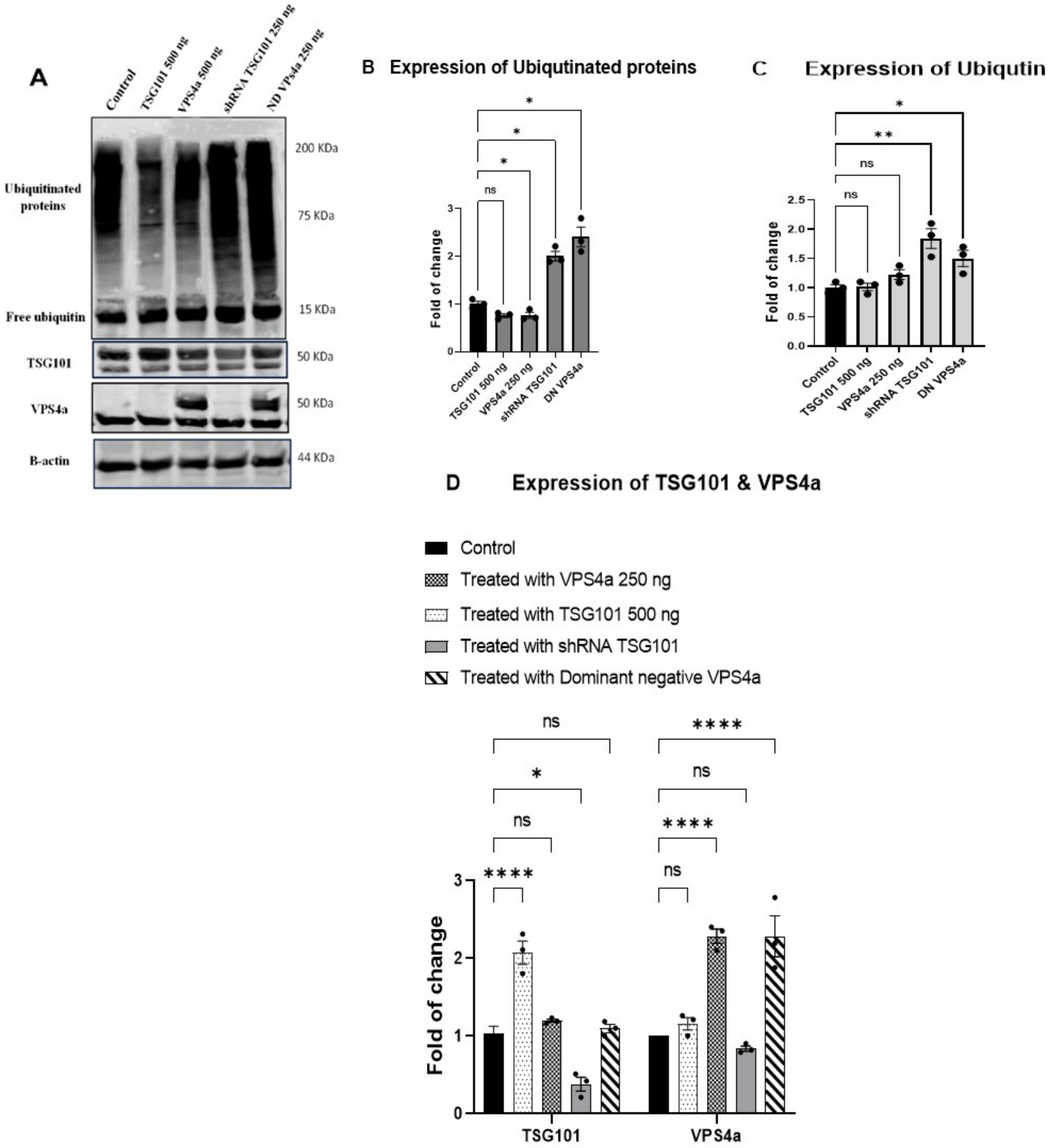


**Fig. S2** Validation of ESCRT modulation and effects on ubiquitin profiles

(A) Representative immunoblots showing expression of ubiquitinated proteins, free ubiquitin, TSG101, VPS4a and β-actin in HEK293 cells under the indicated conditions: Control, TSG101 overexpression (500 ng), VPS4a overexpression (250 ng), shRNA targeting TSG101 (250 ng), and dominant-negative (DN) VPS4a (250 ng). High-molecular-weight ubiquitinated protein species (smear) and free ubiquitin (~15 kDa) are indicated. TSG101 and VPS4a immunoblots confirm effective modulation of ESCRT components. β-actin was used as a loading control.

(B) Quantification of total ubiquitinated protein levels (high-molecular-weight smear) normalised to β-actin and expressed as fold change relative to control. ESCRT perturbation by shRNA TSG101 and DN-VPS4a significantly increased accumulation of ubiquitinated proteins compared with control. Data are presented as mean ± SEM ($n = 3$ independent experiments). Statistical analysis was performed using one-way ANOVA followed by appropriate post hoc testing; *$p < 0.05$; ns, not significant.

(C) Quantification of free ubiquitin levels (~15 kDa band) normalised to β-actin and expressed as fold change relative to control. Modulation of ESCRT components resulted in condition-dependent changes in free ubiquitin abundance. Data are presented as mean ± SEM ($n = 3$ independent experiments). One-way ANOVA with post hoc comparisons; *$p < 0.05$, **$p < 0.01$; ns, not significant.

(D) Quantification of TSG101 and VPS4a protein expression levels under the indicated experimental conditions, normalised to β-actin and expressed as fold change relative to control. Overexpression and knockdown conditions produced the expected alterations in ESCRT component abundance. Data are presented as mean ± SEM ($n = 3$ independent experiments). One-way ANOVA with post hoc testing; *$p < 0.05$, ****$p < 0.0001$; ns, not significant.

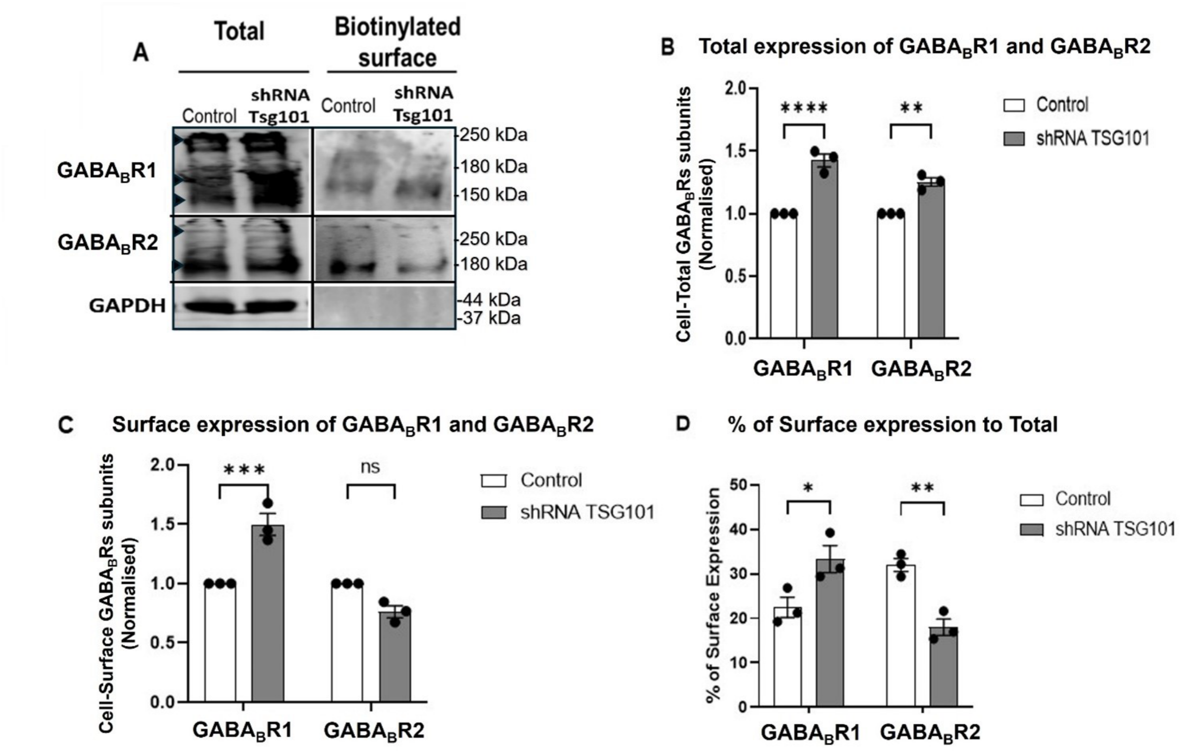


**Fig. S3** TSG101 Differentially Regulates GABA$_B$ Receptor Subunits GABA$_B$R1 and GABA$_B$R2 Through Distinct Trafficking Mechanisms.

Figure A shows representative Western blot images of total and biotinylated surface fractions of GABA$_B$ receptor subunits GABA$_B$R1 and GABA$_B$R2 in HEK293T cells following TSG101 knockdown. GAPDH (44 kDa) served as a loading control and confirmed purity of surface fractions. Densitometric analysis (Figure B) revealed a significant increase in total protein levels for both GABA$_B$R1 (****$p < 0.0001$) and GABA$_B$R2 (**$p < 0.01$), suggesting disrupted turnover or elevated synthesis. However, surface expression patterns diverged: Figure C demonstrates that surface GABA$_B$R1 levels increased significantly (***$p < 0.001$), while GABA$_B$R2 showed no marked change relative to control. These results were further clarified in Figure D, where the surface-to-total ratio revealed opposing trends— GABA$_B$R1 displayed an increased percentage of surface localisation, indicating enhanced trafficking efficiency, whereas GABA$_B$R2 showed a reduced ratio, implying impaired membrane delivery despite elevated total levels. Collectively, these findings indicate that TSG101 exerts receptor-specific regulatory effects on GABA$_B$ subunit trafficking, promoting surface expression of GABA$_B$R1 while disrupting efficient trafficking of GABA$_B$R2. Data are expressed as mean ± SEM. Asterisks indicate statistically significant differences compared to control conditions (*$p < 0.05$, **$p < 0.01$, ***$p < 0.001$, ****$p < 0.0001$).

**Comparative analysis reveals receptor- and ESCRT component-specific patterns of regulation**

| RECEPTOR CLASS | RECEPTOR SUBUNIT | SURFACE EXPRESSION (plasma membrane) | | TOTAL EXPRESSION (whole cell) | |
|---|---|---|---|---|---|
| | | TSG101 KD (ESCRT-I) | DN-VPS4a (ESCRT-III) | TSG101 KD (ESCRT-I) | DN-VPS4a (ESCRT-III) |
| NMDA Receptors | GluN1 | ↑↑ (Strong increase) | ↑ (Increase) | ↑ (Mild increase) | ↑ (Mild increase) |
| | GluN2A | ↑↑ (Strong increase) | ↑ (Increase) | ↑ (Mild increase) | ↑ (Mild increase) |
| AMPA Receptors | GluA1 | ↑↑ (Strong increase) | ↑ (Increase) | ↑/~ (Variable/Minimal) | ↑/~ (Variable/Minimal) |
| | GluA2 | ~ (No/Minimal change) | ↑/~ (Minimal increase) | ~ (No change) | ~ (No change) |
| Kainate Receptors | GluK2 | ↑↑ (Strong increase) | ↑↑ (Strong increase) | ↑ (Mild increase) | ↑ (Mild increase) |
| GABA$_B$ Receptors | GABA$_B$R1 | ↑ (Increase) | ~ (No/Minimal change) | ↑/~ (Variable/Minimal) | ~ (No change) |
| | GABA$_B$R2 | ~ (No/Minimal change) | ~ (No/Minimal change) | ~ (No change) | ~ (No change) |
| GABA$_A$ Receptors | GABA$_A$RG2 | ↑ (Increase) | ↑ (Increase) | ↑/~ (Variable/Minimal) | ↑/~ (Variable/Minimal) |

**CHANGE IN EXPRESSION**

↑↑ Strong increase (50-100% or more) ↑ Increase (~20-50%) ↑/~ Mild increase or variable (~10-20%) ~ No change or minimal (<10%)

**Fig. S4** Comparative analysis of receptor surface and total expression following ESCRT perturbation in primary neurons.

Primary cortical or hippocampal neurons were subjected to TSG101 knockdown (ESCRT-I disruption) or dominant-negative VPS4a expression (ESCRT-III disruption). Surface expression (plasma membrane-localized receptors) and total expression (whole-cell lysates) were quantified for the indicated endogenous neurotransmitter receptor subunits. Data are presented as a heatmap with color intensity reflecting the magnitude of change relative to control conditions (scramble shRNA or wild-type VPS4a). Strong increase (≥50–100%), Increase (20–50%), Mild increase or variable (10–20%), No change or minimal (<10%). NMDA receptor subunits (GluN1, GluN2A) showed robust surface increases following both perturbations. AMPA receptors exhibited subunit-specific patterns (GluA1 > GluA2). GABA receptors displayed heterogeneous responses, with GABA$_A$RG2 and GABA$_B$R1 showing increases following TSG101 knockdown, whereas GABA$_B$R2 was minimally affected. Note that changes in surface expression were not uniformly proportional to changes in total receptor levels, indicating ESCRT-dependent regulation of receptor trafficking and membrane delivery in neurons (created using Biorender).
